# Mosquito excreta reveals circulation of West Nile virus and its underlying ecosystem

**DOI:** 10.1101/2021.12.05.471258

**Authors:** Grégory L’Ambert, Mathieu Gendrot, Sébastien Briolant, Agnès Nguyen, Sylvain Pages, Laurent Bosio, Vincent Palomo, Nicolas Gomez, Nicolas Benoit, Hélène Savini, Bruno Pradines, Guillaume André Durand, Isabelle Leparc-Goffart, Gilda Grard, Albin Fontaine

## Abstract

Emerging and endemic mosquito-borne viruses can be difficult to detect and monitor because they often cause asymptomatic infections in human or vertebrate animals or cause nonspecific febrile illness with a short recovery waiting period. Cases’ detection in vertebrate hosts can be complemented by entomological surveillance, but this method is not adapted to low infection rates in mosquito populations that typically occur in low or non-endemic areas. We identified West Nile Virus circulation in Camargue, a wetland area in South of France, using a cost effective innovative xenomonitoring method based on the molecular detection of virus in excreta from trapped mosquitoes. We also succeeded at identifying the mosquito community diversity dynamic on several sampling sites, together with the vertebrate hosts on which they fed prior to be captured using amplicon-based metagenomic on mosquito excreta without processing any mosquito. Mosquito excreta-based virus surveillance can be considered as a cost-effective and non-invasive strategy that offers the additional asset to reveal the ecological network underlying arbovirus circulation.

## Main

West Nile viruses (WNVs) are mosquito-borne flaviviruses (from the *Flaviviridae* family, *Flavivirus* genus and belonging to the Japanese encephalitis (JE) serogroup) that circulate through several lineages into complex transmission cycles involving mosquitoes from the *Culex* genus and several vertebrate host species, with wild birds as amplifying hosts^1, 2^. WNVs can cause severe and potentially fatal neurological disease in vertebrate hosts, including birds, *Equidae* and humans^3^. Native to Africa, WNVs from lineage 1 were first introduced in Europe presumably through migratory birds in the 1960s^4^ where they caused mainly occasional self-limited outbreaks and sporadic cases^5^ essentially restricted to rural areas. Major European outbreaks occurred in Camargue in 1962-1963^6^ and Romania in 1996^7^. The emergence of lineage 2 in southeastern Hungary in 2004 was associated with a significant upsurge of human and animal cases in several European countries. The largest outbreak was recorded in 2018 with 11 countries reporting 1,548 locally WNV infections, a number that exceeded the cumulative number of all reported infections between 2010 and 2017^8^. WNV lineage 1 and 2 are now circulating in endemic cycles in Europe^5^, causing disease incidence in human every year that tend to be increasingly associated to more urbanized environment. Ecological interactions among bird species that led to WNVs introduction, amplification, and maintenance in local reservoirs, a prerequisite before the virus emergence in humans, are not yet fully understood.

Standard entomological surveillance provides the opportunity to monitor arboviruses circulation in their enzootic cycles based on the processing of thousands of mosquitoes that is cost prohibitive and time consuming. This surveillance method also poses logistical constraints and is thus not adapted to low infection rates in mosquito populations that typically occur in low or non-endemic areas. It has been demonstrated that mosquitoes infected with different arboviruses can excrete large amount of virus genomic RNA^9–11^. Here, we implemented an innovative entomological surveillance strategy inspired from environmental DNA (eDNA) sequencing and molecular xenomonitoring (MX)^12^ that drastically decrease drawbacks associated with standard entomological surveys while providing additional benefits by screening excreta from trapped mosquitoes. We developed a 3D printed lodging that can be adapted on main standard mosquito traps, which offer shelter, food, and a toilet room for trapped mosquitoes. Our xenomonitoring strategy succeeded at detecting two WNV in excreta from the mosquito fauna sampled over a two-month longitudinal study in summer 2020 in Camargue, a wetland area (Rhône delta) in South of France. One of these viruses phylogenetically grouped with a virus from the lineage 1 isolated in 2015 in a horse in the same region. Trapped mosquito excreta were also collected in Mali. Using a single amplicon-based metagenomic approach, we succeeded to monitor the mosquito fauna diversity across different sampling sites and times without processing any mosquitoes. Importantly, we also succeed at identifying vertebrate hosts on which mosquitoes fed prior to be captured. Altogether, we showed that the use of mosquito excreta is efficient to trace arbovirus circulation at low cost and can give precious clues to characterize the ecosystem underlying arbovirus emergence.

## Results

### Virus screening in mosquito excreta

A total of 86 excreta samples were analyzed in this study (80 excreta samples from Camargue, France and 6 from Gao, Mali). These excreta were collected at the end of summer 2020 from trapped mosquitoes contained into the 3D printed MX adapter that has been specifically designed to increase trapped mosquitoes’ survival and to facilitate the recovery of their excreta over several days (Figure 1). A total of 6,845 mosquitoes were recovered from traps in Camargue, comprising 2,819 *Culex spp.*, 3,922 *Ochlerotatus/Aedes spp.*, 91 *Anopheles spp.* and 13 *Culiseta spp.* mosquitoes (Supplementary file 1). *Anopheles spp.* were morphologically identified as *An. hyrcanus sensu lato* and as species from the *An. maculipennis* complex (which includes *An. atroparvus*). Specimens not belonging to the *Culicidae* family were also collected in traps but were not considered in this study. Trapped fauna from Mali were not brought back to Marseille laboratory for analysis. WNV genome was detected by one-step reverse transcription quantitative polymerase chain reaction (RT-qPCR) on excreta samples from 2 traps (2.3%) in Camargue, collected on location F the 15^th^ of September 2020 (cycle threshold (Ct) of 31) and on location G the 22^nd^ of September 2020 (cycle threshold (Ct) of 33) (Figure 2). No excreta samples were found positive for WNV in Mali, nor Usutu virus (USUV) in both Camargue and Mali. Excreta samples from Mali were also tested negative for *Plasmodium .spp.,* the causative agents of malaria, in Mali.

**Figure 1:**
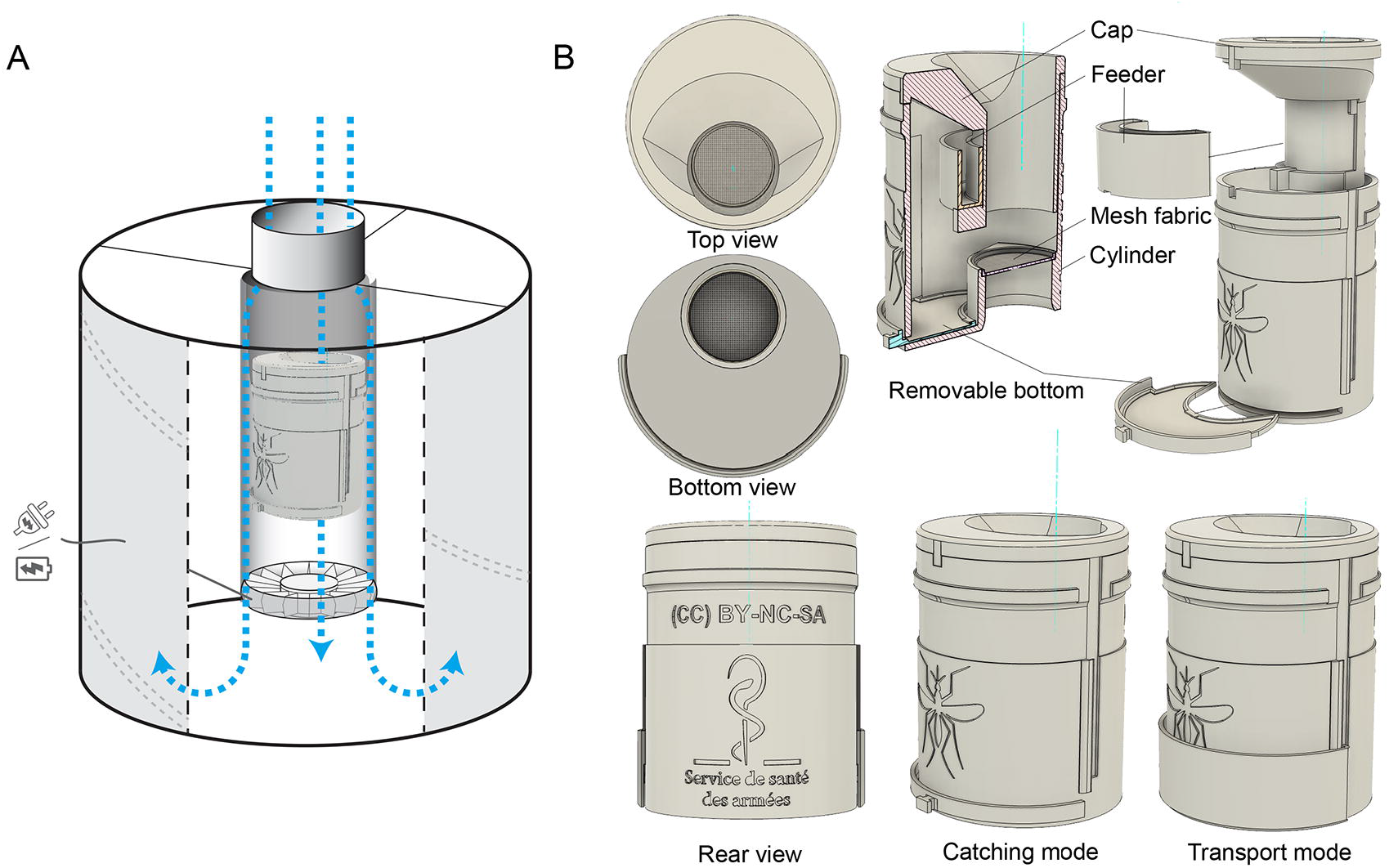
3D representations of the MX adapter designed to increase trapped mosquitoes’ longevity and to collect their excreta for a virus surveillance purpose. (A) Schematic of the MX adapter inside the BG sentinel trap (BGS) in catching mode. (B) Different views of the adapter MX. All components are visible in the cross-sectional and exploded views of the MX adapter: the cap, the mosquito feeder, the ring covered with mesh fabric, the cylinder, and the removable bottom on which can be placed a filter paper to collect mosquito excreta. In transport mode, the sliding gate covers the open slot in absence of the removable bottom. The MX adapter was created on Fusion 360 (AutoDesk). MX adapter is under the Creative Commons (CC) license BY-NC-SA (Licensees may copy, distribute, display and make derivatives only for non-commercial purposes and by giving credits to the authors). All printable files are provided in Supplementary file 3.

**Figure 2:**
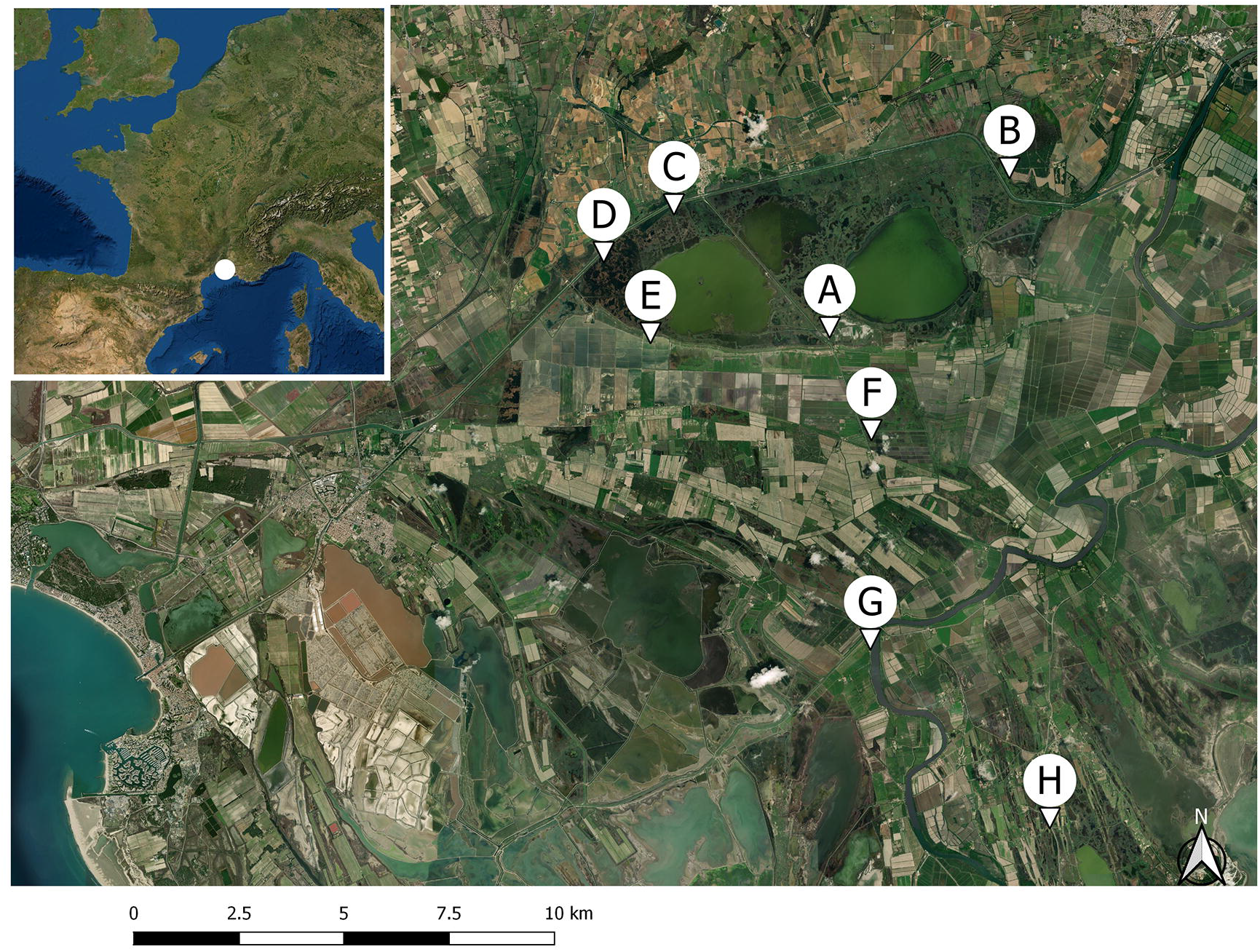
Study map with the geo-localization of the 8 sampling sites (A to H) in the Camargue wetland in South of France. The Map was created using the Free and Open Source QGIS Geographic Information System using satellite imagery from the Environmental Systems Research Institute (ESRI).

In Camargue wetland, a total of 1 *Culex*, 17 *Ochlerotatus /Aedes* and 1 *Anopheles* mosquitoes were collected in the first positive trap (F 2020/09/15) while 5 Culex, 31 *Ochlerotatus/Aedes* and 1 *Anopheles* mosquitoes were identified in the second trap (G 2020/09/22). No WNV viruses were detected in these mosquitoes by RT-qPCR after an individual extraction. The amount of excreta recovered on these traps was higher than it would be expected by considering that they would be expelled by these mosquitoes only, suggesting that a mosquito escape issue could have occurred for these traps. *Culex* and *Culiseta* mosquitoes from traps tested as negative for WNV and USUV based on excreta, were screened for these viruses based on an individual grinding and a pooled extraction and detection procedure. These viruses were not detected in any of the 241 pools, each made of 12 mosquitoes.

We succeeded at sequencing a 341 bp WNV genomic region (GenBank accession number OK489805) located on the envelope gene on one excreta sample that was tested positive by RT-qPCR with the lowest Ct (sample F 2020/09/15). Phylogenetic analysis revealed that the WNV identified in our study belonged to lineage 1 and was closely genetically related to a WNV isolated on a symptomatic horse the 3^rd^ of October 2015^13^ (GenBank accession number MT863559) (Figure 3 A). Our WNV small genomic section diverged from the 2015 WNV (GenBank accession number MT863559) by 5 single nucleotide polymorphism (SNPs) and from a WNV isolated in Spain on a horse in 2010^14^ (GenBank accession number JF719069) by 4 SNP (Figure 3 B).

**Figure 3:**
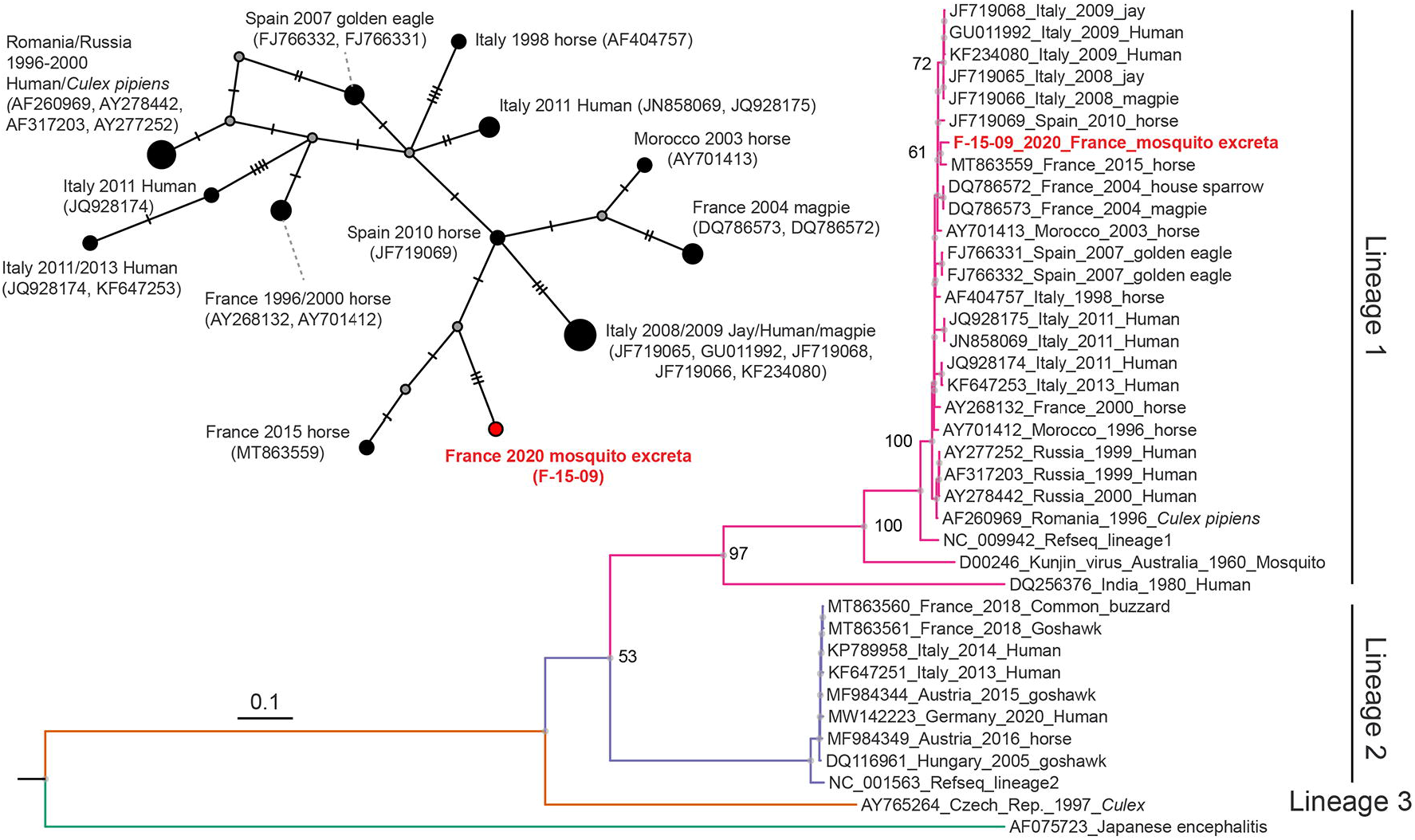
Molecular relationship at the intraspecific level of WNV genomes from lineages 1, 2 and 3, with the virus identified in this study in mosquito excreta. (A) Molecular phylogenetic tree of 38 WNV genome, including the 341 bp genome section identified in this work. The phylogenetic analysis was performed on a curated alignment of 10,302 bp by allowing gap positions within the final block. One sequence of JEV (AF075723) was used as an outgroup in the phylogenetic tree. The evolutionary history was inferred using a maximum likelihood method using PhyML. Bootstrap values obtained after 100 replicates are shown at major nodes (percentage of replicate trees in which the associated taxa clustered together). Evolutionary distances, as represented by length of branches, are expressed in number of base substitutions per site. (B) Haplotype network inferred by the TCS method using the same sequences as above trimmed to the length of our sequenced amplicon (341 bp). The size of each circle represents the frequencies of the haplotype. Mutations are shown as perpendicular bars along the branches and grey small circles represent inferred unsampled haplotypes. The WNV genome section identified in this study is indicated in red.

### Amplicon-based metagenomic analysis on trapped mosquito excreta

The mitochondrial DNA mixture extracted from mosquito excreta samples was amplified over a ∼ 460 bp mitochondrial DNA section corresponding to a sub fragment of the classical Folmer cytochrome c oxidase subunit I (COI) fragment^15^ routinely used in metabarcoding analysis. The sequencing generated a total of 3,968,028 reads across all samples with a mean of 45,610 (1^st^ quartile: 40,335; 3^rd^ quartile: 50,797) reads per sample, including the control. All paired-end reads were turned into chimera-free amplicon sequence variants (ASVs) after a trimming, quality-based filtering, merging, dereplicating and denoising steps^16^. The total number of reads was reduced to 7,752 ASVs with a median abundance of 28,290 (1st quartile: 23,735; 3rd quartile: 33,305) per sample. Taxonomy was next assigned to each ASV using *blast* on a database gathering Fungi, Protist, and animal COI records. A total of 1,330 (17.6%), 1,681 (22.3%), and 53 (0.7%) ASVs were assigned to the *Animalia*, *Fungi*, and *Protist* kingdoms, respectively, and 4,486 ASVs (59.4%) were unassigned. Unassigned ASVs corresponded to off-targeted genomic amplifications or assignments falling below the required 80% identity threshold. ASVs count per kingdom represented 17% (N=406,892), 39% (N= 924,375), 0.1% (N=2,292) and 44% (N= 1,027,285) of the total ASVs count for the *Animalia*, *Fungi*, *Protist* kingdoms and unassigned ASVs, respectively (Supplementary file 2). Fungi from the genera *Leohumicola* (class: *Leotiomycetes*, phylum: *Ascomycota*), *Tremella*, *Cryptococcus* and *Rhodotorula* (phylum: *Basidiomycota*) were overrepresented. The *Arthropoda* phylum was the most represented inside the *Animalia* kingdom (96% of ASVs abundance) with most ASV counts assigned to the *Insecta* class.

### Analysis of the Culicidae diversity based on trapped mosquito excreta

Both *Arachnida* and *Insecta* classes were represented in the pool of DNA recovered inside the adapter MX. The *Diptera* order represented 78% of ASVs abundance inside the *Insecta* class and taxon from the *Culicidae* family represented 48% of ASVs counts inside the *Diptera* order (Supplementary file 2). This was consistent with the fact that mosquitoes (*Culicidae*) were not the only specimen recovered from the traps. One hundred and three ASVs could be assigned at the species level for 14 mosquito species, *i.e. Ochlerotatus detritus* (6 ASVs), *Ochlerotatus caspius* (1 ASV), *Culiseta subochrea* (3 ASVs), *Culex theileri* (8 ASVs), *Cx. quinquefasciatus* (1 ASV), *Cx. pipiens* (13 ASVs), *Cx. modestus* (30 ASVs), *Anopheles Hyrcanus* (12 ASVs), *Anopheles atroparvus* (2 ASVs), *An. gambiae sensu lato* (1 ASV), *Aedes vexans* (5 ASVs), *Ae. albopictus* (3 ASVs), and *Ae. aegypti* (18 ASVs). Thirty-eight, 4 and 1 ASVs were assigned to the *Ochlerotatus*, *Culex* and *Uranotaenia* genera, respectively, and 31 were assigned to the *Culicidae* family (Figure 4-A).

**Figure 4:**
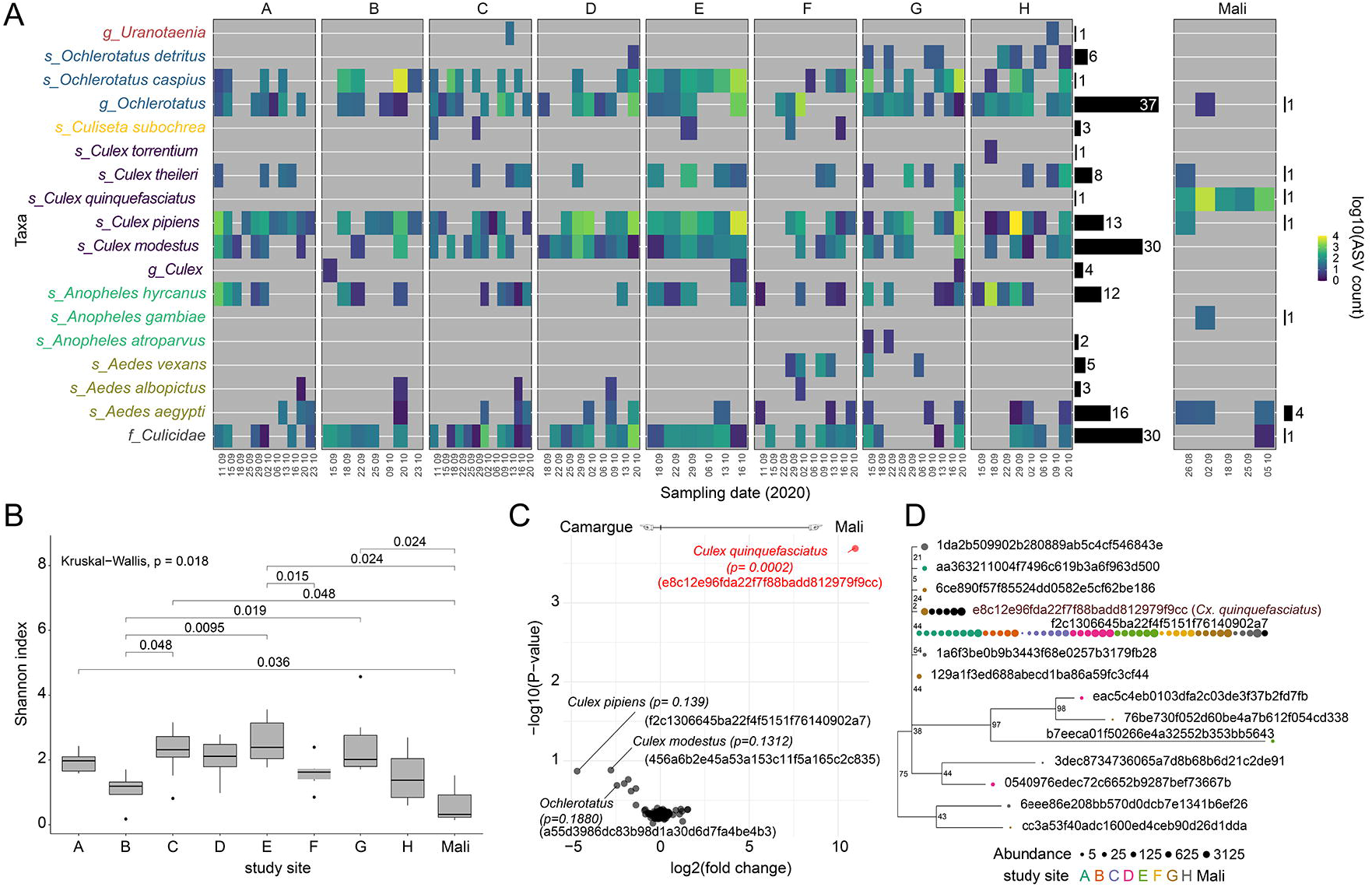
Amplicon-based metagenomic analysis of the *Culicidae* diversity through the sequencing of trapped mosquito excreta. A- Heatmap representing the total number of ASV (log10 scale) attributed to a taxon according to sampling sites and sampling times. The number of ASVs assigned to each taxon is represented with horizontal black bars at the right- hand side of the graph for Camargue and Mali study sites. B-*Alpha* diversity (Shannon’s diversity index) across study sites. Kruskal-Wallis test was used to compare Shannon’s diversity across sample sites. Non-adjusted *p* values are presented. C- Volcano plot representing the magnitude of change (X axis) versus the -log10 expected *p* value of Welch’s t test. Red represents features called as significantly differentially abundant with *p* < 0.05. D- Phylogenetic relationship among ASVs assigned to *Culex pipiens s.l.* mosquitoes. The phylogenetic tree was derived from ultrafast bootstrap algorithm implemented in Qimme2 to calculate phylogenetic distances among ASVs. Study sites in which the ASV was identified are represented with a color code.

Shannon index was used to quantify the species diversity and richness inside each sample (*Alpha* diversity). *Alpha* diversity varied from 0.66 ± 0.75 (mean ± SD, Shannon index) for the Mali study site to 2.57 ± 0.73 for site E in Camargue and was not significantly different across sites when considering a Holm or Bonferroni *p*-value correction (Figure 4-B). The *Alpha* diversity in samples from Mali was significantly lower than the *Alpha* diversity observed on all other sites in Camargue (2 ± 0.83, all site from Camargue confounded, *p* = 0.01). Significant *Alpha* diversity difference was observed over time (*p* = 0.03, mixed-linear regression).

The *Culicidae* compositional differences among samples (*Beta* diversity) were significantly affected by the study area (*i.e*. Camargue vs Mali) with all nominal pairwise comparison between Mali and Camargue sites being under the significant threshold (*p*-value < 0.05, PERMANOVA on weighted UniFrac distances), with the exception of site H (*p* = 0.158). *Beta* diversity was also significatively different between site D and sites B, C, F and G in Camargue (Supplementary file 4). *Culicidae* species composition in site B was also significantly different from site A and E, and between site C and H. None of these pairwise comparisons were significant when considering Benjamini-Hochberg FDR correction to adjust the *p*-values to account for the multiple testing. The ASV assigned to the *Cx. quinquefasciatus* species (feature ID: e8c12e96fda22f7f88badd812979f9cc) was significatively associated to community composition differences between Camargue and Mali, accounting for the differential abundance of ASVs between these study areas (Aldex2, *p*=0.0002, Expected *p* value of Welch’s t test, Figure 4-C). The abundance of an ASV assigned to the mosquito species *Cx. modestus* (feature ID: 456a6b2e45a53a153c11f5a165c2c835, Supplementary file 4) was significantly over- represented in sites D and E in all their pairwise comparisons described above (*p* < 0.05). At the qualitative level (*i.e*., ASV presence/absence), ASVs assigned to *An. gambiae* species were only detected in Mali, as expected. *An. atroparvus* was only detected in site G in Camargue and *Ae. vexans* only in sites F and G. ASVs assigned to *Ae. albopictus* and *Ae. aegypti* were detected in Camargue. These species were not identified in trapped mosquitoes in this study area and are not likely to occur in Camargue, suggesting spurious ASV assignments for these species.

Amplicon-based genomic was further used to assess the genetic diversity inside the *Cx. pipiens s.l.* mosquito species complex. We could reveal several *Culex* cytochrome oxidase c subunit I (COI) haplotypes circulating in Camargue (Figure 4-D). The haplotype assigned to *Cx. quinquefasciatus* was over-represented in the Mali study site but do not branch apart on the tree topology (Figure 4-D). ASVs genetic diversity was not obviously associated to *Cx. pipiens pipiens* and *Cx. pipiens molestus* cryptic forms inside the *Cx. pipiens s.l.* complex, as revealed by the tree topology incorporating representative sequences from all forms (Supplementary file 5).

Mosquito species composition revealed on DNA present in mosquito excreta (amplicon-based metagenomic) was compared with the species composition revealed by morphological identification of trapped mosquitoes at the genus level for all sample collections from Camargue. The *Aedes* and *Ochlerotatus* genera were grouped for simplicity in this analysis. Mosquito species composition shared 68% similarity between both identification methods when considering the symmetrical Sokal & Michener similarity index, with 111 (33.6%) and 113 (34.2%) matches for the presence revealed by both methods and absence revealed by both methods, respectively. A total of 44 (13%) and 62 (18.8%) mismatches was revealed for the presence in metagenomic/absence in trapped mosquitoes and absence in metagenomic/presence in trapped mosquitoes, respectively. The similarity dropped to 51% when using the asymmetric Jaccard index that do not consider species absences in both methods. Importantly, Jaccard similarity index was much lower when considering under- represented mosquito species, *i.e. Anopheles* (Jaccard index: 20%) or *Culiseta* (Jaccard index: 7%) genera, as compared to the highly represented *Aedes/Ochlerotatus* (Jaccard index: 58%) or *Culex* (Jaccard index: 73%) genera.

### Chordata diversity revealed by amplicon-based metagenomic based on trapped mosquito excreta

The *Chordata* phylum represented 4% of ASVs abundance inside the *Animalia* kingdom, with most ASVs assigned to the *Homo sapiens* species (77% of *Chordata*), followed by species from the *Artiodactyla* order (*Suidae* (pigs) and *Bovidae*, 11% of *Chordata*) and birds (6% of *Chordata*). At the qualitative level, *Homo sapiens*, *Rattus norvegicus*, bats from the *Vespertilionidae* family and pigs (*Sus scrofa*) species were identified in both Camargue and Mali study sites. ASVs assigned to a frog species (*Hoplobatrachus occipitalis*) and the *Tayassu pecari* species were uniquely identified in Mali. Mali is inside the frog species distribution area but the geographic distribution of *pecari* is in South America, suggesting a spurious identification for this species. Six fish’s species were also detected in this study. ASVs assigned to these fish’s species (class: *Actinopterygii*) were scarce (3% of the *Chordata* phylum), and *Salmo salar* (salmon) was the most represented and present in 3 study sites in both Camargue and Mali.

Vertebrates typically associated to the Camargue regions were detected, such as *Bos taurus*, *Equus caballus* or birds belonging to the *Ardea* genus (*i.e.* herons and egret). While accounting for only 0.3% of ASVs counts inside the *Animalia* kingdom, ASVs assigned to the *Aves* class were the most diverse inside the *Chordata* phylum with 6 birds identified as the species level (*i.e. Sylvia atricapilla*, *Motacilla flava*, *Hirundo rustica, Pica pica*, and *Corvus corax* from the *Passeriforms* order, and *Gallus gallus)*. ASVs were also assigned to the *Ardea* genus, *Passeriformes* order and *Phasianidae* family (Figure 5). *Alpha* diversity at the *Chordata* level varied from 0.60 ± 0.37 (mean ± SD, Shannon index) for study site E to 1.28 ± 0.76 for site B in Camargue and was not significantly different across sites but varied significantly over time (*p* = 0.03, mixed-linear regression).

**Figure 5:**
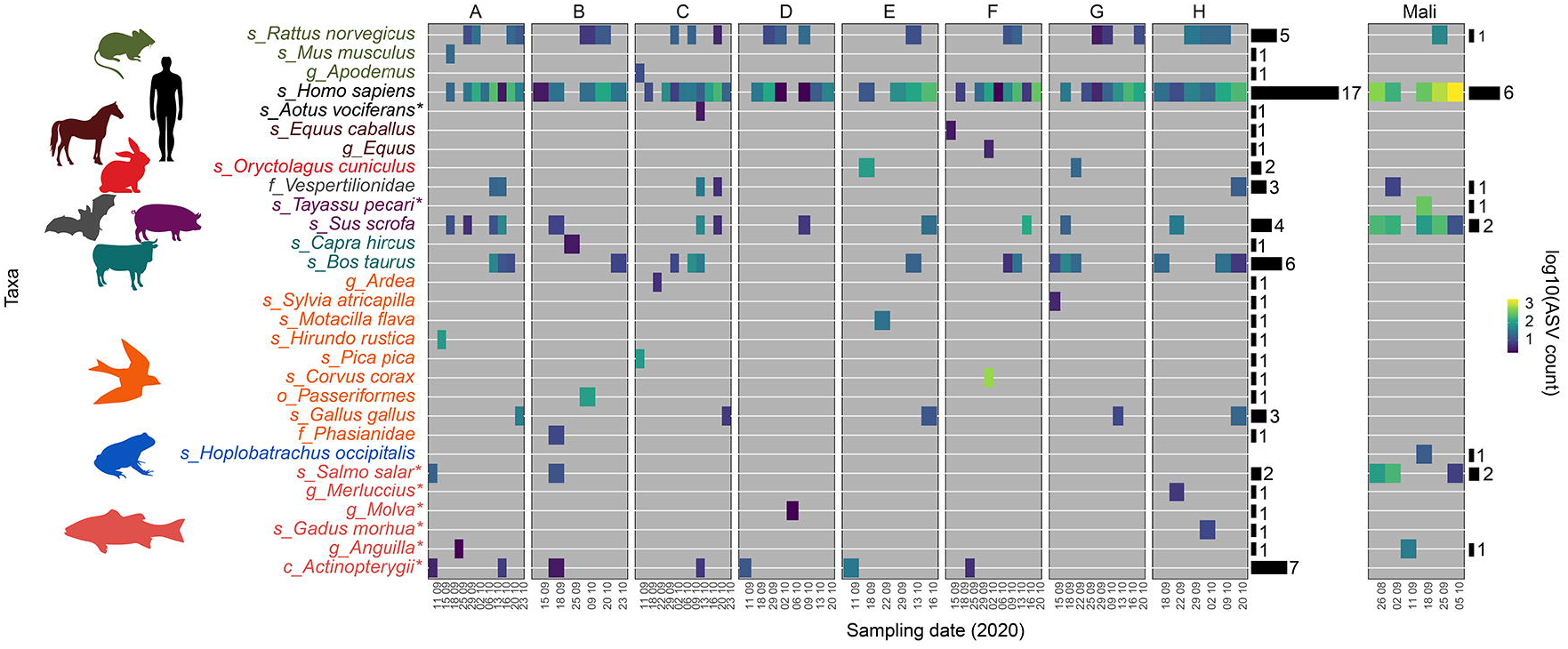
Vertebrate (*Chordata*) diversity identified based on digested blood through the sequencing of trapped mosquito excreta. Heatmap representing the total number of ASV (log10 scale) attributed to a taxon according to sampling sites and sampling times. The number of ASVs assigned to each taxon is represented with horizontal black bars at the right-hand side of the graph for Camargue and Mali study sites.

## Discussion

WNV surveillance is usually achieved through the monitoring of symptomatic cases in human and animals^17, 18^, seroprevalence studies^19–21^, dead birds^2, 22^, blood donor^23, 24^ or virus screening in trapped mosquitoes^25^. High rates of asymptomatic arbovirus infections, the broad immunological cross-reactivity across flaviviruses, the difficulties to report and collect dead birds^26, 27^, and low arbovirus infection prevalence in mosquitoes are hindering the use of these methods to early-detect arbovirus circulation with an accurate dating of infection events. The discovery that mosquitoes with a systemic arbovirus infection can massively excrete virus RNA^9, 11^ has open new perspectives in entomological surveillance of viruses^10, 28, 29^. Indeed, virus identification in trapped mosquito excreta considerably alleviates the processing charge and costs associated with entomological surveillance by extending the period between mosquito trap collections and making optional the steps of mosquito species identification, pooling, and testing. This non-invasive method that does not require strong entomological knowledge can also be extended to monitor mosquito-borne parasites^30, 31^.

Here, we succeeded at identifying WNV circulation in Camargue over a short 6-week period without the need to process any mosquito. This is only the third time that WNV have been detected in mosquitoes in this region since 196^13, 32, 33^ despite important efforts to search for infected vectors, including after epidemics. The sequencing of a short genomic region on the envelope gene indicated its affiliation to WNVs from lineage 1 (Western Mediterranean clade) that circulated in Southern Europe in the last 20 years^13^. These viruses can be maintained locally from one year to another in enzootic cycles that involve resident bird species and ornithophilic mosquitoes by overwintering in hibernating infected adult mosquitoes^34, 35^ or in bird species with chronic infection^36, 37^. Another hypothesis would be recurrent introductions through migratory birds that would carry the viruses northward during spring migration from African wintering places where these viruses are endemic, or from West Europe in late summer when birds return to Africa. Implementing active WNV surveillance in paths and resting places of migratory birds in Europe and Africa would help to resolve the mechanisms of WNV maintenance or resurgence in Europe. Unfortunately, no infected mosquitoes were recovered from the traps positive for WNV based on detection on mosquito excreta due to mosquito escape issues that occurred at the beginning of this study. Complete genome virus sequencing and WNV infection prevalence would have been easier to achieve by processing trapped mosquitoes as a second intention. In case of a mosquito escape due to a trap failure or a faulty handling, arbovirus RNA and trapped mosquito DNA can still be recovered in the excreta with our system. For instance, we could assess the mosquito species diversity and composition in Mali without an access to the trapped mosquitoes. Mosquito excreta collected on filter papers are also easy to transport and store from a remoted field site to the laboratory even in an absence of cold storage.

We succeeded to have a global view of the mosquito species composition and abundance in different sampling sites. Mosquito species emblematic to Camargue such as *Ochlerotatus caspius* and *Culex pipiens* were identified via their DNA contained in their excreta in Camargue but not in Mali. Taxonomic identification is limited with our method to the genetic diversity that can be captured and amplify over a short mitochondrial sequence. Here, taxonomic identifications are biased toward arthropods because we used a degenerated primer set specifically designed to amplify arthropods^38, 39^. The use of primers targeting vertebrates’ mitochondrial DNA on the same target might further increase our ability to identify the diversity of vertebrate species on which mosquitoes fed prior to be trapped^40^. These limitations might have explained the spurious or unexpected species identifications in this study. *Aedes albopictus* and *Ae. aegypti* COI sequences might share a strong homology on our target region with other *Aedes* species in Camargue that are not present in our database. The identification of fish species in this study might have resulted from any DNA contamination from the environment (*e.g*. food contamination), laboratory reagents or kits^41^. ASVs assigned to these fish species were scarce and filtering out low-abundant and/or low-prevalent features might improve false-identifications, with the trade-off of losing species identification. The *Cx. pipiens s.l*. complex has two recognized forms - *pipiens* and *molestus* – that exhibit behavioral and physiological differences and can form hybrids. *Culex pipiens pipiens* feed preferentially on birds, and *Cx. pipiens molestus* on humans but both can transmit WNV^42, 43^. Our method could reveal genetic diversity inside the species level but could not associate *Cx. pipiens s.l*. COI haplotypes to one of these biotypes. Failure to discriminate these forms inside the complex using mitochondrial DNA has been previously described^44, 45^. Additional nuclear genomic regions could be targeted in the metagenomic scheme to further discriminate these species inside the complex^43^ or to non-invasively screen for known molecular signatures (*e.g.* insecticide resistance).

Potential WNV bird reservoir species were also non-invasively identified on trapped mosquito excreta. Determining the reservoir role of local bird species for WNV is not straightforward due to the challenging task of obtaining blood samples from animal for diagnoses purposes. It usually involves wild bird captures that are difficult to operationalize or blood-meal analysis of trapped mosquitoes prior to implement experimental infection studies^46, 47^. Blood-fed mosquitoes can be accidentally captured in CO2 baited traps or be trapped during their quest to complete their blood-meals. It is also possible to specifically target blood-engorged mosquitoes in their resting sites, but this method implies an active prospection and is difficult to implement for exophilic species. Our method captures vertebrate blood as it is incrementally digested by trapped mosquitoes without the need to process mosquitoes during their digestion stage. Blackcaps (*Sylvia atricapilla*), Barn swallows (*Hirundo rustica*), Yellow wagtails (*Motacilla flava*) and herons or egrets identified in this study were previously listed as bird species potentially involved in the introduction, amplification and spread of WNV in Camargue^48^. They are all bird species present in Camargue during the mosquito season that migrate in West Africa during winter. WNV antibodies have been detected in most of these species^49^. WNV has also been isolated from common magpies (*Pica pica*)^50^ in 2004 in Camargue. *Corvidae* are sedentary birds that are potential hosts to amplify the virus and to bring it closer to the urban areas. Our method cannot directly link a virus collected in trapped mosquitoes to a vertebrate host. However, it can reveal trophic preferences and the range of potential hosts involved in WNV introduction and amplification in different areas. Ecological factors associated with WNV outbreaks are not yet fully understood. Increased host diversity can either reduce risk of virus emergence by diluting the abundance of a vector or a reservoir host inside a community^51^ or increase the risk if the abundance of a vector is a function of vertebrate host diversity^52^. Extending the entomological survey over several diverse locations and seasons might help to assess if bird or mosquito species diversity and composition can influence the risk of WNV emergence.

Costs and logistic constraints are impeding the implementation of nation-wide entomological surveillance programs worldwide. Here, we used both the RNA and DNA genomic materials contained into excreta from trapped mosquitoes to (i) non-invasively survey the emergence and spread of WNV, (ii) monitor mosquito species diversity without the processing of mosquitoes and (iii) to invade ecological network underlying WNV emergence by identifying vertebrate animals on which trapped mosquitoes fed before to be captured, using a single and easy to implement amplicon-based metagenomic procedure. Our cost-effective strategy can be considered as a complementary early-warning tool to detect the circulation of mosquito-borne pathogens affecting human health or to non-invasively survey the presence of these pathogens in remote areas without health-care system.

## Methods

### 3D-printed BG sentinel mosquito trap modification

We created a 3D-printed adapter (MX adapter) to increase mosquito longevity in BG Sentinel (BGS, Biogents AG, Regensburg, Germany) traps, inspired from the device designed by Timmins D.R. and colleagues^53^. Briefly, our system is composed of a 120-mm high and 87/90 mm (inside/outside) diameter cylinder with a removable cap that is inserted into the depressurized BGS catch pipe. The MX adapter is attached beneath the intake funnel and is replacing the original catch bag. The original catch bag was cut at the bottom to house the MX adapter (Figure 1A). The MX adapter is traversed longitudinally by an airflow piping to allow air depressurization inside the catch pipe, and thereby allowing aspiration at the top of the BGS intake funnel via the fan located underneath the system. The airflow piping is cut with its upward pointing section sealed with mesh fabric to prevent mosquito escape from the device while letting the air flows through. A mosquito feeder, filled with a cotton ball soaked in 10% sugar water, is attached to the inner side of the cylinder. The MX adapter has a removal bottom on which a filter paper (Whatman, grade 3, ref. 1003-917) can be placed when the system is set in mosquito catching mode. A vertical sliding gate can be used to seal the system during transport when the bottom has been removed (optional) (Figure 1B). The MX adapter provide a safe and moisturized enclosure for trapped mosquito with an easy access to sugar. Mosquito excreta can be easily collected on the filter paper at the bottom of the adapter. First version of BGS traps were used in this study but our device can be adapted to BG-Sentinel 2 and CDC light traps (Supplementary figure 1). The MX adapter was created on Fusion 360 (AutoDesk) and 3D printed in either PLA or PETG on a Sigmax R19 (BCN 3D). MX adapter 3D files (.stl format) are provided in supplementary file 3 under the Creative Commons (CC) license BY-NC-SA.

### Study area and samples collection

The study was carried out in a ∼130 km² area in Camargue (Figure 2), a large wetland in the Southeast of France with a temperate Mediterranean climate located inside the Rhone river delta. The Camargue is a nature reserve that hosts large populations of resident and migratory birds and is famous for its white horses and black bulls. West Nile virus transmission has been repeatedly reported in Camargue for several years^50, 54, 55^. BGS traps were placed at the ground level at 8 sites nearby the “Marais du Charnier”, each separated by few kilometers. Adult mosquitoes were captured over 6 weeks from the 11^th^ of September 2020 to the 23^rd^ of October 2020, at the end of the mosquito season, using BGS modified with the MX adapter, as described previously. Mosquito captures were conducted over 3 to 4 consecutive days, with traps collection and reconditioning for a new mosquito trapping session occurring twice a week. Carbon dioxide was provided as a mosquito attractant for every BGS traps. Pressurized carbon dioxide bottles and 12V power packs were used to operate over time. During trap collection, the MX adapter was removed from the BGS and maintained 5 minutes on a sealed plastic box filled with dry ice to knock down mosquitoes. Knock down mosquitoes were then transferred to 50 mL tube and transported alive to the laboratory. This mosquito collection procedure avoids mosquitoes to escape from the device and was implemented late in our mosquito collection session (after the end of September) in response to mosquito escape issues coming from the fields. Filter paper covered with mosquito excreta were placed into annotated sealed plastic bags and transported at ambient temperature to the laboratory. Once in the laboratory, mosquitoes and filter papers were stored at -20°C until the extraction procedure. Adult mosquitoes were identified morphologically in the laboratory at the genus level and sorted by genus, trap, and collection date. One trap was implemented in Gao, Mali, from the 26th of August 2020 to the 5th of October 2020 in continuous operation with 6 sampling over this period. No mosquito was brought back to the lab and only mosquito excreta were analyzed for these samples.

### RNA/DNA extraction

#### RNA/DNA extraction from filter papers with mosquito excreta

Virus detection was first conducted on mosquito excreta collected on filter papers. Briefly, filter papers were coiled and placed individually at the bottom of 14 mL plastic tubes (Falcon, ref: 352059) before to be soaked in 3 mL of Lysis buffer RAV1 (NucleoSpin 96 virus core kit, Macherey⃞Nagel, Düren, Germany) for 5 minutes. Ten microliters of phage MS2 (RNA) were added to each tube as an internal extraction control. Filter papers were then manually grinded with a 5 mL pipette. Several 1 mm diameter holes were drilled in the transparent caps of the 14 ml tubes to create a colander, and then clipped on each tube. Closed tubes were placed upside-down on a larger 50 mL tube (with the colander cap placed downward, at the bottom of the 50 mL tube) and centrifuged 5 min at 3,000 rpm. Flowthroughs were collected and mixed with 3 mL 96–100% ethanol before to be loaded on NucleoSpin Virus Columns in several steps. RNA/DNA extraction was performed as indicated by the manufacturer indications. Eluates were kept at -20°C until use.

#### RNA/DNA extraction from captured mosquitoes

Virus detection in large number of samples is time consuming and cost prohibitive, especially when low infection rates are expected in mosquito populations. Here, we implemented a strategy based on the simultaneous grinding of 96 individual mosquitoes in a 96-well plate inspired from Holleley C. and Sutcliffe A. MR4 protocol^56^, followed by an extraction and detection in pool of 12 mosquitoes (one lane of a plate). This strategy drastically reduces labor and costs associated to virus screening, as compared to virus detection in individual mosquitoes, while still providing the opportunity to sequence or isolate virus in single mosquito in a second intention, and to obtain accurate prevalence estimates. Briefly, *Culex* mosquitoes were placed individually in each well of a shallow conical bottom 96-well plate with 80 µ L of buffer composed of 1.3 mL of Proteinase K resuspended in proteinase buffer according to the manufacturer instructions (NucleoSpin 96 virus core kit, Macherey-Nagel) in 9 mL PBS. A male *Ae. albopictus* infected with the insect specific *Aedes flavivirus* (AEFV) was add on well position H12 of each plate as an internal extraction control. All mosquitoes from a plate were simultaneously grinded for 5 minutes by moving up and down and rocking vigorously a disposable bacterial colony replicator tool. Plates were then incubated at 70°C for 1 hour with the bacterial colony replicator tool left on the top of the plate. After a short grinding step post-proteinase K digestion, the replicator tool was discarded and 15 µL of each well from a same plate line were pooled. Plates with individually grinded mosquitoes were stored at -20°C. 500 µL of AVL lysis buffer (QIAamp Viral RNA kit, Qiagen, Hilden, Germany) was added to each pool with 5 µL of phage MS2 as an additional internal extraction control. After 10 minutes of incubation at room temperature, 500 µ L of 100% ethanol was added to each pool and RNA was extracted according to the manufacturer protocol (QIAamp Viral RNA kit, Qiagen).

### Viruses and *Plasmodium spp.* detection

Detection of WNV and USUV genomic RNA was performed with a one-step reverse transcription quantitative polymerase chain reaction (RT-qPCR) assay. The multiplexed RT-qPCR was performed using the SuperScript one-step RT-PCR Platinum® Taq Mastermix (Invitrogen, Cergy Pontoise, France), MS2 primers and probe (21347398), or either WNV or USUV specific primers and probe (Supplementary table 1) using 10 µL of buffer 2x, 0.8 µL of forward and reverse primers, 0.3 µL of probe, 0.8 µL of enzyme and 2.3 µ L of RNase-free water for one reaction, with the following cycling protocol : 48°C for 30 min, 95°C for 10 min, 50 cycles at 95°C for 15 sec, 60°C for 1 min). Amplification was performed on a CFX real-time thermocycler (Bio-Rad, Hercules, CA, USA). Concerning the detection of AEFV in infected male *Ae. albopictus* internal controls (extraction procedure for trapped mosquitoes in pools), virus genomic RNA was first reverse transcribed to complementary DNA (cDNA) with random hexamers using M-MLV Reverse Transcriptase (Life Technologies, Inc., Gaithersburg, Maryland, USA) according to the manufacturer’s instructions. cDNA was amplified by 35 cycles of PCR using the corresponding set of primers described in Supplementary table 1 and amplicons were visualized by electrophoresis on a 1% agarose gel. *Plasmodium spp.* detection was performed using 10 µL of 2x SYBR Green PCR Master Mix (Thermofisher scientific, Waltham, USA), 0.5 µL of forward and reverse 16 µM primers (Supplementary table 1), 4 µL of RNase-free water and 5 µL of DNA extract for one reaction using the following cycling protocol: 95°C for 5 min, 45 cycles at 95°C for 15 sec, 60°C for 1 min. Amplification was performed on a LightCycler (Roche Diagnostics, Mannheim, Germany).

### Virus sequencing

We sequenced WNV genome based on the tiling amplicon-based sequencing (PrimalSeq) method developed by Quick J.^57^, and Grubaugh N.D.^58^ and colleagues. Briefly, PrimalSeq protocol generates overlapping amplicons of ∼⃞400 base pairs from 2 multiplexed PCR reactions to generate sufficient templates for subsequent high-throughput sequencing. We used the multiplex primer scheme developed for WNV lineage 1 that have been validated by Grubaugh N.D. *et.al.*^58^. To identify the WNV lineage before whole genome sequencing, we sequenced in first intention 5 overlapping genomic sections covering the envelop gene with a set of primers degenerated to amplify WNV lineages 1 and 2. Illumina Nextera® universal tails sequences were added to the 5’ end of each of these primers to facilitate the library preparation by a two-step PCR approach (WNV lin.1 & 2, Supplementary table 1). Purified RNAs from excreta samples positive for WNV in Q-RT-PCR were first reverse transcribed to complementary DNAs (cDNAs) with random hexamers using M-MLV Reverse Transcriptase (Life Technologies) according to the manufacturer’s instructions. For the multiplex PCR reactions, 5 μl of cDNA was used in a 20 μl reaction mixture made of 5 μl of Hot START 5X BIOamp DNA Polymerase mix (Biofidal, Lyon, France), 4 μl of forward and reverse primer mix at 10 μ and 11 μl of water. The thermal program was: 10 min of polymerase activation at 96°C followed by 35 cycles of (i) 30 sec denaturing at 96°C, (ii) 30 sec annealing at 62°C and (iii)1 min extension at 72°C, followed by a final incubation step at 72°C for 7 min to complete synthesis of all PCR products. A 15 cycles PCR was then performed using Nextera® Index Kit – PCR primers, that adds the P5 and P7 termini that bind to the flow cell and the dual 8 bp index tags. Resulting amplicons were purified with magnetic beads (SPRIselect, Beckman Coulter), quantified by fluorometric quantification (QuantiFluor® dsDNA System, Promega) and visualized on QIAxcel Capillary Electrophoresis System (Qiagen). Libraries were sequenced on a MiSeq run (Illumina) using MiSeq v3 chemistry with 300bp paired-end sequencing.

After demultiplexing, trimmomatic v0.33^59^ was used to discard reads shorter than 32 nucleotides, filter out Illumina adaptor sequences, remove leading and trailing low-quality bases and trim reads when the average quality per base dropped below 15 on a 4-base-wide sliding window. Reads were aligned to the WNV lineage 1 reference genome (RefSeq entry: NC_009942) with bowtie2 v.2.1.0^60^. The alignment file was converted, sorted and indexed using Samtools v0.1.19 . Coverage and sequencing depth were assessed using bedtools v2.17.0^62^. The consensus sequence was generated with Ivar^58^ with default options.

### Phylogenetic analyses

A background set of 36 full-length WNV genome sequences across different lineages was obtained from GenBank. One full-length genome sequence of Japanese Encephalitis Virus was also downloaded to be used as an outgroup in the phylogenetic tree. Genome sequences were aligned using Clustal Omega and curated by Gblocks software implemented in the seaview v.5.0.4 interface^63^ by allowing gap positions within the final block and less strict flanking positions before to generate the best-scoring maximum-likelihood (ML) tree with 100 bootstrap replicates with PhyML (15980534). The GTR + I + G nucleotide substitution model was chosen based on the lowest Akaike Information Criterion (AIC) value using the Smart Model Selection in PhyML (SMS) software^64^. Phylogenetic trees were visualized using the ggtree R package^65^. All sequences were trimmed to the length of our sequenced amplicon (341 bp) before to be imported into the PopArt program^66^ to create a TCS haplotype network^67^.

### Amplicon-based metagenomic

The amplicon-based metagenomic method implemented here was performed on mosquito excreta eluates post NucleoSpin Virus RNA/DNA extraction. Even though the NucleoSpin Virus Columns kit is optimized to extract virus RNA/DNA, the eluate also contained non- viral RNA/DNA. Virus detection, sequencing and metagenomic approaches were performed on the same eluates to facilitate the procedure. A ∼ 460 bp mitochondrial DNA section corresponding to a sub fragment of the classical Folmer cytochrome c oxidase subunit I (COI) fragments^15^, was amplified with universal primers BF2/BR2^68^ (Supplementary table 1) that are routinely used for macroinvertebrate monitoring^38, 39^. Illumina Nextera® universal tails sequences were added to the 5’ end of each of these primers to facilitate library preparation, as described above for the WNV tiling amplicon-based sequencing procedure. PCR amplifications, library preparation and sequencing were made according to the same protocol described above for WNV sequencing excepting for the annealing temperature that was set to 50°C for 30 sec. All raw sequences have been deposited in the NCBI database under the NCBI BioProject number PRJNA768434.

### Taxonomic identification and diversity indices

Demultiplexed fastq sequences were imported to QIIME2 version 2021.2 for bioinformatics analyses. The qiime2-dada2 pipeline^16^ was used for turning paired-end fastq files into merged reads, filtering out Illumina adapters, denoising and removal of chimeras and filtering out replicates. Taxonomic assignment was carried out for the amplicon sequence variants (ASVs) using the qiime2-feature-classifier classify-consensus-blast plugin using a database of 1 176 764 sequences gathering Fungi, Protist, and Animal COI records, recovered from the Barcode of Life Database Systems (BOLD) the 7^th^ March 2021. The database was built according to the step-by-step tutorial provided by Devon O’Rourke^69^. Only sequences within our primer coordinates were retained in the final database. A percentage identity threshold of 90% was used to assign a taxonomy to an ASV.

Shannon’s diversity index was used to assess *Culicidae* and *Chordata* species richness and evenness (relative abundance of species inside a sample), *i.e. Alpha* diversity, across time and sampling sites. Kruskal-Wallis test was used to compare Shannon’s diversity across sample sites using the compare_means function implemented in the ggpubr R package^70^. Bonferroni *p*-*value* correction was used to account for the multiplicity of tests. A linear mixed-effects model implemented with the lme4 R package^71^ was used to assess differences in *Alpha* diversity across time. A random effect was implemented on the study site. Data were rarified to 150 (*Culicidae* analysis) and 50 (*Chordata* analysis) reads per sample, before to assess the compositional differences among samples (*Beta* diversity) using a permutational analysis of variance (PERMANOVA) on weighted UniFrac distances that compare species compositions considering both the relative abundance and phylogenetic distances among species. Benjamini-Hochberg FDR correction was used to adjust the *p*-values to account for the multiple testing. Phylogenetic distances between species from the *Culicidae* family and *Chordata* phylum were calculated with IQ-TREE^72^ with the ultrafast bootstrap algorithm and automatic model selection, as implemented in the Qiime2 phylogeny iqtree-ultrafast-bootstrap plugin. Differential ASVs abundances between sampling sites were compared on a non-rarefied dataset using the q2-aldex2 differential abundance package, by modelling the data as a log-ratio transformed probability distribution rather than as counts^73^. The R package phyloseq^74^ was used to represent phylogenetic relationship among ASVs assigned to *Culex pipiens* mosquitoes. All further diversity metrics implemented in Qiime2 can be accessed by running Qiime2 diversity commands on data provided on Supplementary files 4.

Matching coefficient of Sokal & Michener index (symmetrical) and Jaccard index (asymmetrical) were used to compare the *Culicidae* fauna composition revealed based on trapped mosquitoes identified morphologically and amplicon-based metagenomic on the corresponding mosquito excreta for all sample collections from Camargue at the genus level. Considering a contingency table of binary data such as n11 = a, n10 = b, n01 = c and n00 = d, the Sokal & Michener index was computed as follow: s = (a+d) / (a+b+c+d), and Jaccard index as follow: s = a / (a+b+c). The latter is an asymmetrical index that does not treat double zero (d) in the same way as double presences (a) as a reason to consider samples similar.

All statistical analyses were performed in the statistical environment R. Figures were made using the package ggplot2^75^ and the Tidyverse^76^ environment. The Map was created using the Free and Open Source QGIS Geographic Information System using satellite imagery from the Environmental Systems Research Institute (ESRI).

## Acknowledgments

We are grateful to Laurie-Lou Weghel for her interest and support throughout this project. We also thank Frédéric Jean (EID Méditerranée) for his thorough involvement on the field work and Nicolas Benoit for his help regarding the molecular detection of *Plasmodium spp*. We also thank Clément Gendrot and Sylvain Buffet for their help at designing and printing the first version of the adapter MX, it all started from there. We are also thankful to the Qiime2 contributors and community for providing such an easy, enjoyable, and pre-built access to metagenomic analyses.

## Data availability

Amplicon-based metagenomic raw sequencing data are accessible under the NCBI BioProject number PRJNA768434. The WNV genomic section sequenced in this project is accessible under the GenBank accession number OK489805. Qiime2 artifacts with all amplicon-based data are available in Supplementary file 4.

## Supplementary information

**Supplementary table 1:** Molecular amplification systems used in this study. Oligonucleotides sequences (primers and probes) are presented with their corresponding species, gene targets and amplicon sizes. Illumina Nextera® transposase sequences that have been added to primers during the PCR amplification step of the metagenomic method are represented in green.

**Supplementary file 1:** Mosquito species inventory. Number of mosquitoes identified morphologically at the genus level in each trap. *Anopheles* mosquitoes were further identified at the species or species complex level.

**Supplementary file 2: Interactive representation of the metagenome of trapped mosquito excreta displayed with a Krona chart.** The number 2,360,844 represents the total number of ASV counts across all samples. Taxonomy nodes are shown as nested sectors arranged from the top level of the hierarchy at the center and progressing outward. Navigational controls are at the top left, and details of the selected node are at the top right.

**Supplementary file 3:** MX adapter 3D files in .stl format. MX adapter is under the Creative Commons (CC) license BY-NC-SA (Licensees may copy, distribute, display, and make derivatives only for non-commercial purposes and by giving credits to the authors).

**Supplementary file 4:** Archive comprising (i) Qiime2 code that was used in the amplicon-based metagenomic analysis pipeline, (ii) the data base with sequences recovered from public repositories (bold_COI_seqs_ref_db.qza and bold_COI_taxa_ref_db.qza), (iii) alpha-rarefaction analysis (alpha_rarefaction_curves.qzv), (iv) metadata file associated to the data (COI-metadata.txt), (v) denoised features (rep-seqs-trim.qza/.qsv) and their associated taxonomy (taxonomy_blast-trim_80identity.qza/.qsv). PERMANOVA results for *Culicidae* (weighted-unifrac-study_site_significance_Culicidae.qzv) and *Chordata* (weighted-unifrac- study_site_significance_Chordata.qzv) are provided as well as Aldex2 differential abundance analysis between sites from Camargue (Aldex_figures.html). All further diversity metrics implemented in Qiime2 can be accessed by running Qiime2 diversity commands on data provided. All .qsv files can be easily loaded on the Qiime2 visualizer at https://view.qiime2.org.

**Supplementary file 5:** A phylogenetic tree displaying the intra *Cx. pipiens s.l.* complex genetic diversity revealed by our method relative to the *Cx. pipiens* and *Cx. molestus* forms. Haplotypes revealed in our study are represented with their feature ID in dark red. GenBank accession numbers of sequences representative to *Cx. quinquefasciatus*, *Cx. pipiens* form *molestus* and *Cx. pipiens* form *pipiens* are represented in green, blue, and black, respectively. The evolutionary history was inferred using a maximum likelihood method using PhyML. The GTR + I nucleotide substitution model was chosen based on the lowest Akaike Information Criterion (AIC) value using the Smart Model Selection in PhyML (SMS) software. Evolutionary distances, as represented by length of branches, are expressed in number of base substitutions per site.

**Supplementary figure 1:** Picture of the MX adapter fit to a CDC light trap.

