## Supplementary file 2 for "Mosquito excreta reveals circulation of West Nile virus and its underlying ecosystem": Supplementary file 2.html

Javascript must be enabled to view this page.

magnitude
magnitudeUnassigned

feature\_table\_taxonomy\_outputRsum

2360844

6
406892

78429
389841

37

32

30

30

30

2

2

2

5

5

5

5

12453
7440

254
420

49

49

49

115

115

115

2

2

2

49
3319

308

308

308

65

65

65

848

848

848

854

854

854

309

309

309

3
486

483

483

63

63

30

33

6

6

6

322

322

322

9

9

9

76
475

280

280

280

21

21

21

10

10

10

21

3

18

18

3

3

3

64

64

42

42

42

42

757

736

736

736

21

21

21

70

70

70

70

70

4

4

4

4

4

20534
298271

496
28016

22614
20453

2161

349
4906

12

12

4545

4545

1053

1053

1053

1053

4310
5105

4
10

6

6

410
3

407

150
22

128

28

28

28

128

128

128

58
69

11

75

4

4

4

14
71

3

3

54

54

3748
6098

7

7

7

701
729

28

28

7

7

7

415
22

393

393

257
231

24

2

2

935

935

935

2779
233138

46

124

1739

1688

187

1499

2

43

43

8

8

8

8

67
6028

3

5775
2

5486

118

169

124

124

59

379

379

379

26147
22368

1080

1080

83

5

78

2585

2585

13

13

18

18

924
404

10

206

206

304

304

6585

3615

1446

125

2042

2

20

20

2948

2948

2

2

8

8

883

883

42

73

73

73

57
53015

288

288

19195

19195

1225

1225

3951

3951

17279

17279

34

34

329

329

801

801

14

14

147

147

2
619

617

13

13

6

6

8971
2

78

2

79

2156

6654

5

5

25
63

28

6

4

18

18

482

9

9

16
450

421

13

23

7

7

20

20

20

146

6

6

140

140

20990

7

7

20983

20983

28

28

11095
112685

44

57

57

5282

22

370

4890

5960

9

25

5926

14
54259

1547

13985

4

32783

5926

3

3

16221
35985

3663

15939

162

112

112

112

112

1132
174

25

25

25

10

10

10

85

85

85

829

2

827

9

9

9

1015
3008

155

112
155

43

14

14

14

812

812

812

5

5

5

781

781

768

13

9

9

9

217

217

217

577

19

19

19

19

209
13

64

64

64

3
132

64

64

47
65

18

349

16

16

16

23

23

23

310

17045

1047

5

5

5

947
92

27

27

27

4

4

4

80

80

80

744

662

662

82

82

95

95
9

86

86

15506

395

395

11

21

21

363

363

111

111

111

111

7

7

7
4

3

13055

13052

13052

13052

3

3

3

1837

340

340

340

1127

1127

1127

370

3

3

367

367

101

101

15

15

15

15

15

104
477

38

38

38

17

6

6

11

2

9

9

318

318

318

318

1027285

924375
6

526972
14356

2

2

2

2

2

15
122

107

107

13

13

13

13

13

379
12285

11906
4981

11
4

7

404
2637

19

19

2202

12

12

4
12

8

8

4265

3

3

4262

4262

1635
500194

5
2

3

3

10

10

10

10

498544

498544

399532
498544

99005

2

5

397397
2232

8576
2653

32

32

32

32

2227
5863

3

3

3

33

33

3

3

3

114

114

114

20
16

4

4

3433

3433

30

30

13
3

5

5

5

5

5

5

15
4

11

11
6

5

98039

7

7

7

98032

98032

98032

98032

708

708

708

708

708

4

4

4

4

287838

287838

52
287838

236745

236745

51041
48678

219

2144

2292

2292

14
2292

2053

11
2053

87
105

5

13

1937
1903

8

13

9

4

89

89
49

3

37

37

136

133

123

10

10

3

3
