## Supplementary figures and images for "Mosquito excreta reveals circulation of West Nile virus and its underlying ecosystem"

### Supplementary figure 1

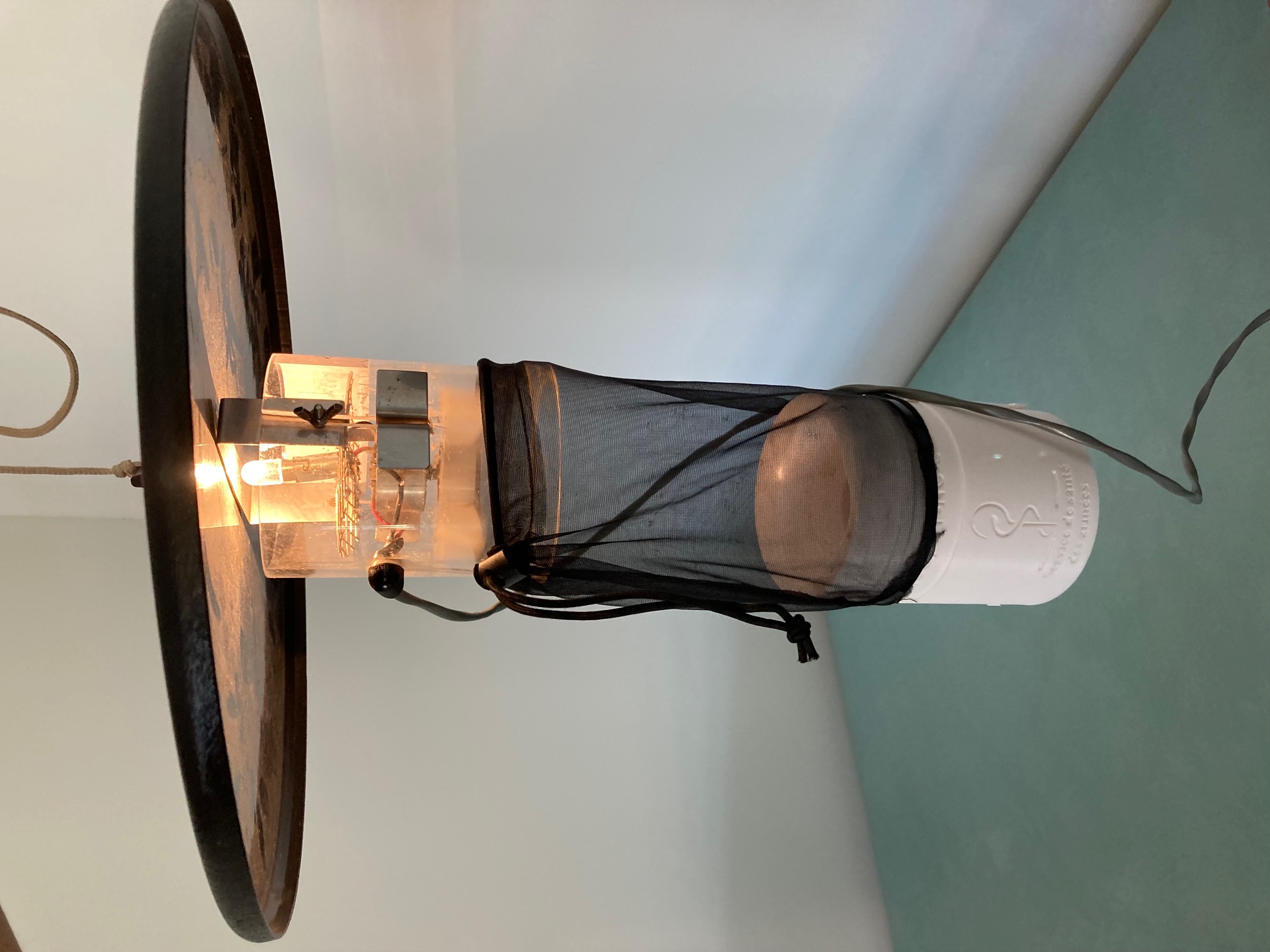

### Supplementary file 5

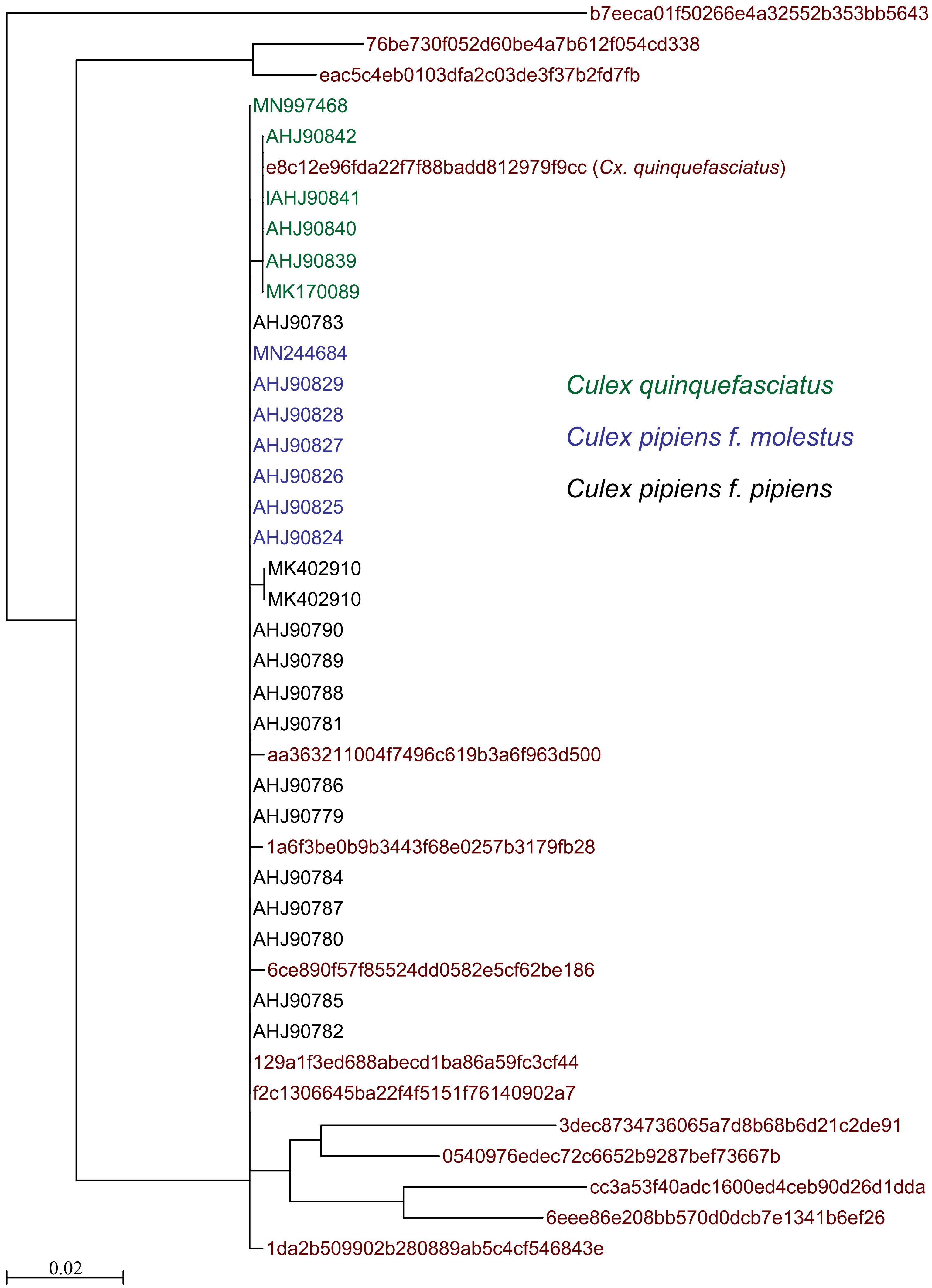
